# Localization of regions of a *Bordetella pertussis* autotransporter, Vag8, interacting with C1 inhibitor

**DOI:** 10.1101/2020.03.09.983296

**Authors:** Naoki Onoda, Yukihiro Hiramatsu, Shihono Teruya, Koichiro Suzuki, Yasuhiko Horiguchi

**Author notes:** Corresponding author: Yukihiro Hiramatsu.

## Abstract

*Bordetella pertussis* is the causative agent of pertussis (whooping cough), a contagious respiratory disease that has recently seen a resurgence despite high vaccination coverage, necessitating improvement of current pertussis vaccines. An autotransporter of *B. pertussis*, virulence-associated gene 8 (Vag8), has been proposed as an additional component to improve pertussis vaccines. Vag8 is known to play a role in evasion of the complement system and activation of the contact system by inactivating the complement regulating factor, C1 inhibitor (C1 Inh), which inhibits serine proteases, such as plasma kallikrein (PK). However, the nature of the molecular interaction between Vag8 and C1 Inh remains to be determined. In the present study, we attempted to determine the minimum region of Vag8 that interacts with C1 Inh by examining the differently–truncated Vag8 derivatives for the ability to bind and inactivate C1 Inh. The region of Vag8 from amino–acid residues 102 to 548 was found to bind C1 Inh and cancel its inhibitory action on the protease activity of PK at the same level as a Vag8 fragment from amino–acid residues 52 to 648 covering the passenger domain, which carries its extracellular function. In contrast, the truncated Vag8 containing amino–acid residues 102 – 479 or 202 – 648 barely interacted with C1 Inh. These results indicated that the two separate regions of amino–acid residues 102 – 202 and 479 – 548 are likely required for the interaction with C1 Inh.

**Importance:** Pertussis is currently reemerging worldwide, and is still one of the greatest disease burdens in infants. *B. pertussis* produces a number of virulence factors, including toxins, adhesins, and autotransporters. One of the autotransporters, Vag8, which binds and inactivates the complement regulator C1 Inh, is considered to contribute to the establishment of *B. pertussis* infection. However, the nature of the interaction between Vag8 and C1 Inh remains to be explored. In this study, we narrowed down the region of Vag8 that interacts with C1 Inh and demonstrated that at least two separate regions of Vag8 are necessary for the interaction with C1 Inh. Our results provide insight into the structure–function relationship of the Vag8 molecule and information to determine its potential role in the pathogenesis of *B. pertussis*.

## Text

*Bordetella pertussis* causes pertussis (whooping cough), a contagious respiratory disease that has recently seen a resurgence despite high vaccination coverage (1, 2), which has prompted attempts to improve current pertussis vaccines. A number of groups have attempted to identify novel bacterial components that confer efficient immunity against *B. pertussis* infection, and proposed virulence-associated gene 8 (Vag8) as a possible candidate (3, 4). Vag8 is an autotransporter, which is autonomously secreted by an intramolecular system consisting of passenger and translocator domains. After passing through the inner membrane by a canonical secretion system, the passenger domain is translocated across the outer membrane by the translocator domain. The passenger domain of Vag8 is cleaved and liberated into the extracellular milieu (5). In *B. pertussis* infection, the liberated Vag8 binds and inactivates the complement regulating factor, C1 inhibitor (C1 Inh), which inhibits serine proteases involved in the complement system and the plasma kallikrein (PK)-kinin system (6), and plays a role in evasion of the complement system and activation of the contact system by the bacterium (5, 7, 8). However, the nature of the molecular interaction between Vag8 and C1 Inh remains unknown. Here, we attempted to localize the minimum region of Vag8 that interacts with C1 Inh by examining different truncated Vag8 derivatives for the ability to bind and inactivate C1 Inh. Our results suggest that at least two separate regions of Vag8 are necessary for interaction with C1 Inh.

### Binding of truncated Vag8 derivatives to C1 Inh

We generated 9 types of truncated Vag8 derivatives together with the wild-type, Vag8_WT_, covering the passenger domain (Fig. 1A). The mobility of each recombinant protein on SDS-PAGE followed by immunoblotting corresponded to that estimated from its molecular mass (Fig. 1B). The ability of each recombinant Vag8 to bind C1 Inh was quantified by ELISA-based binding assay. Among the C-terminal deletion mutants, Vag8_52-596_ and Vag8_52-548_ bound C1 Inh to similar extents to Vag8_WT_, while the binding of Vag8_52-479_ was markedly reduced (Fig. 2A, left panel). The N-terminally deleted derivatives showed decreased binding levels with extension of deletion (Fig. 2A, center panel). Both-terminal deletion derivatives, Vag8_102-596_ and Vag8_102-548_, bound C1 Inh similarly to Vag8_WT_, while the binding of Vag8_102-479_ was markedly diminished (Fig. 2A, right panel). Taken together, these observations indicated that the region from amino–acid residues 102 to 548 is a prerequisite for binding to C1 Inh.

**FIG 1.**
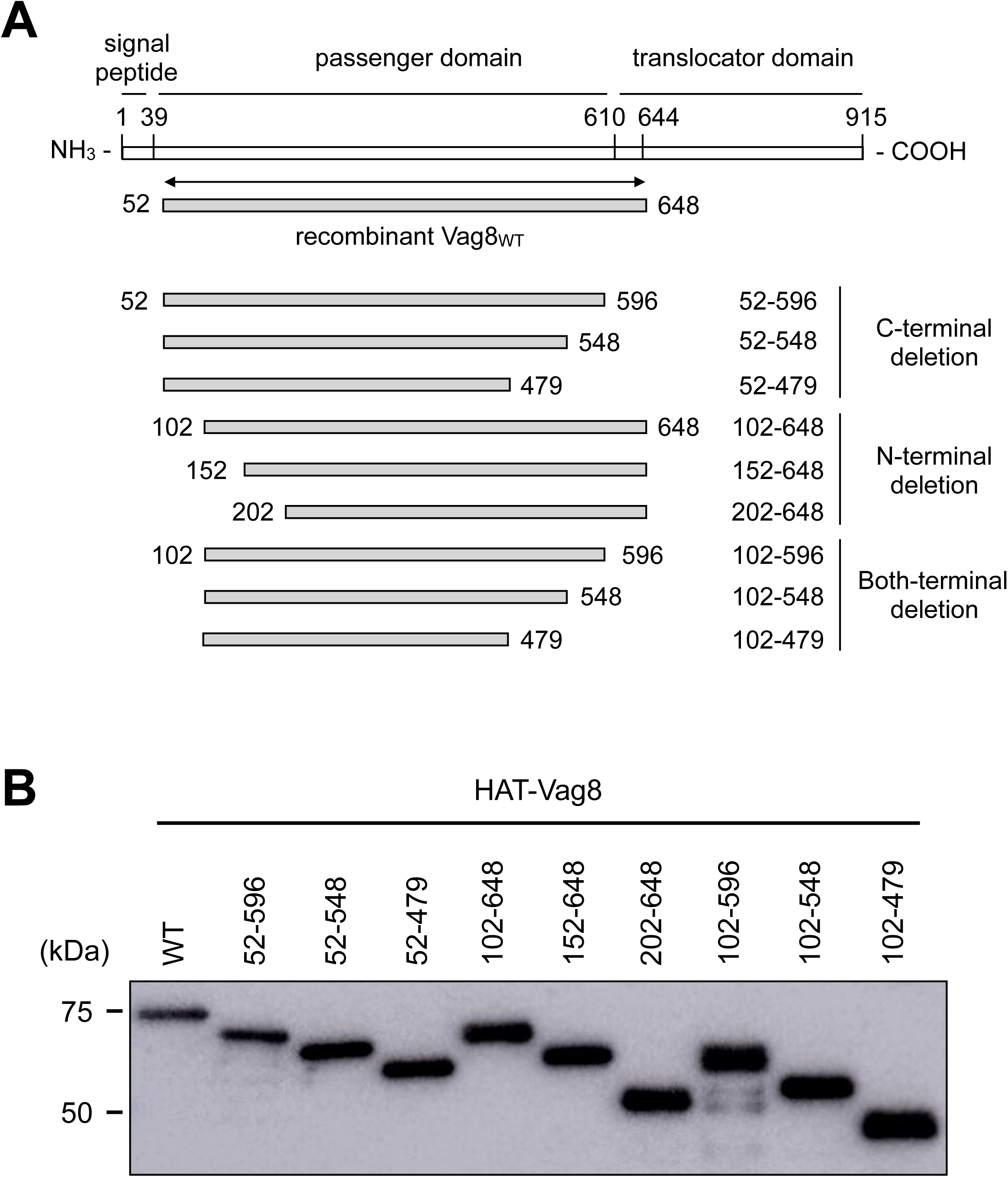
Vag8 and its derivatives used in this study. (A) Schematic representations of Vag8 and each recombinant protein are shown. Vag8_WT_ and truncated derivatives of Vag8 are listed with their names and corresponding amino–acid positions. Each recombinant protein has a HAT-tag at the N-terminus. (B) Immunoblotting of the recombinant Vag8_WT_ and truncated derivatives.

**FIG 2.**
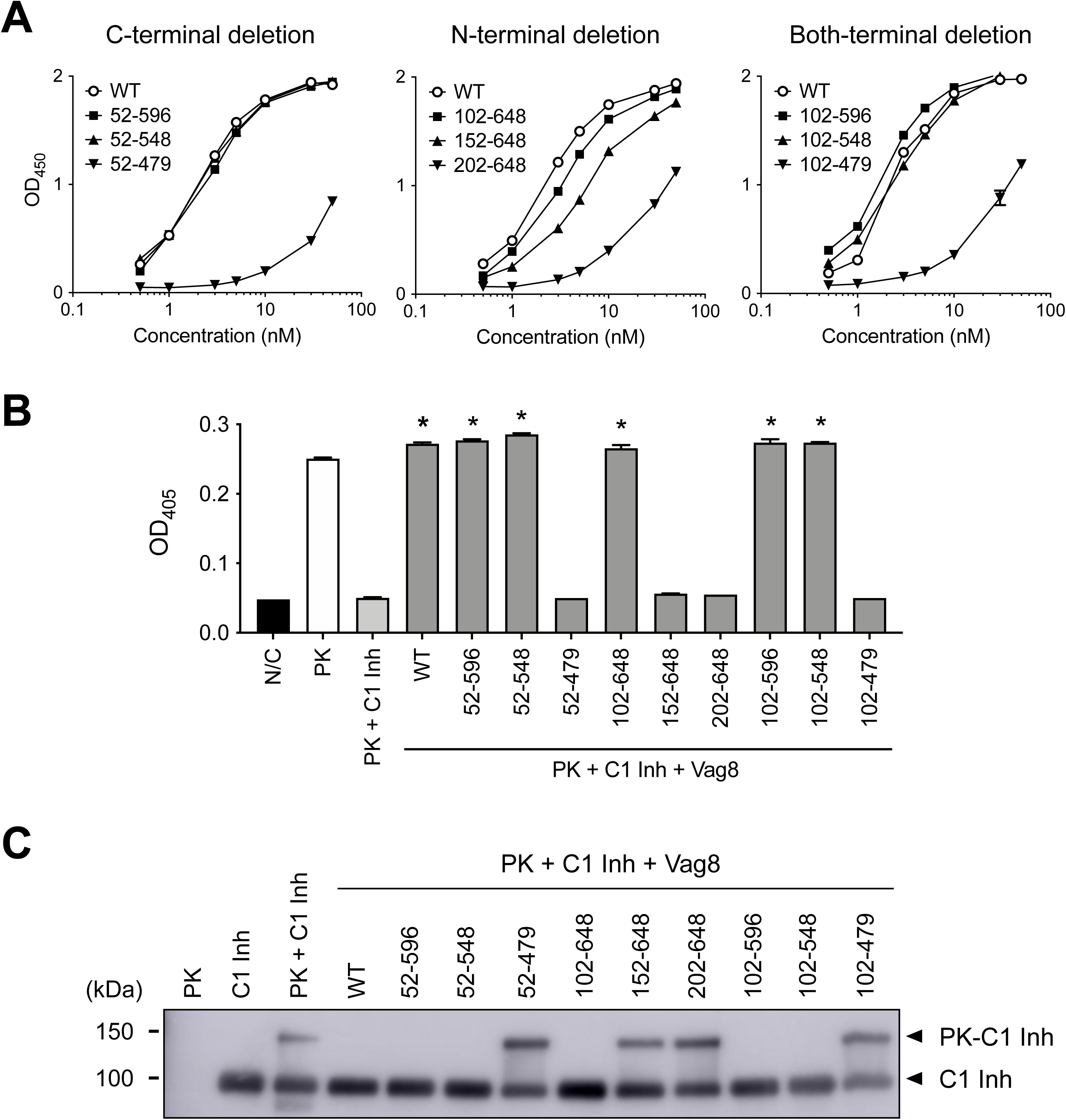
Abilities of Vag8 and its derivatives to interact with C1 Inh. (A) Binding of truncated Vag8 derivatives to C1 Inh analyzed by ELISA-based binding assay. The data are presented as means (n = 3) and SEM. (B) Cancellation of the inhibitory effects of C1 Inh on PK activity by truncated Vag8 derivatives. N/C is the negative control without PK, C1 Inh, and Vag8s. The data are presented as means (n = 3) and SEM. Statistical analyses were carried out by one-way analysis of variance and Tukey’s multiple comparison test using Prism 8 (GraphPad Software). \**P* < 0.01. (C) Immunoblotting of C1 Inh incubated with PK and truncated Vag8 derivatives. Note that C1 Inh bound covalently to PK to form a complex with larger molecular mass (upper arrowhead) (10).

### Cancellation of inhibitory effects of C1 Inh by Vag8s

The interactions between C1 Inh and the Vag8 derivatives were also examined based on the inhibitory effect of C1 Inh on the protease activity of PK (7, 9). Vag8_52-596_, Vag8_52-548_, Vag8_102-648_, Vag8_102-596_, and Vag8_102-548_ inhibited the C1 Inh activity to the same level as Vag8_WT_, while Vag8_52-479_, Vag8_202-648_, and Vag8_102-479_, which scarcely bound to C1 Inh, did not (Fig. 2B). Vag8_152-648_, which showed intermediate binding to C1 Inh, did not inhibit the C1 Inh function. In addition, we examined whether the truncated Vag8 derivatives inhibit formation of the complex of PK and C1 Inh, which was detected by mobility shifts on SDS-PAGE followed by immunoblotting. The truncated Vag8 derivatives carrying the minimum region for association with C1 Inh inhibited covalent binding between PK and C1 Inh, while the others did not (Fig. 2C).

## Conclusions

We demonstrated that the region of Vag8 consisting of amino–acid residues 102 – 548 retains the ability to bind C1 Inh and to cancel its inhibitory action on PK. As neither Vag8_102-479_ nor Vag8_202-648_ showed the full activity of intact Vag8, the regions of amino–acid residues 102 – 202 and 479 – 548 are likely required for the interaction with C1 Inh, suggesting that at least two separate regions of Vag8 mediate the interaction with C1 Inh. Although further studies are required, the present study provide insight into the mechanism by which Vag8 inhibits C1 Inh activity.

## Methods

### (i) Recombinant Vag8 derivatives

DNA fragments encoding WT and truncated derivatives of Vag8 were amplified from *B. pertussis* 18323 using appropriate primers (Table S1 in the supplemental material), and cloned into the *Xho*I-*Eco*RI or *Xho*I-*Hin*dIII sites of pCold II-HAT (3). The expression of each Vag8 derivative from *Escherichia coli* BL21 (DE3) or DH5α harboring each expression vector was induced with 1 mM isopropyl β-D-1-thiogalactopyranoside. The collected bacteria were lysed in 50 mM sodium phosphate buffer, pH 8.0, containing 300 mM NaCl (Buffer A) by sonication. After centrifugation, the pellets were dissolved in Buffer A containing 8 M urea and dialyzed against Buffer A. The resultant samples were centrifuged, and the supernatants were independently applied to a column of HIS-Select Nickel Affinity Gel (Sigma-Aldrich) equilibrated with 50 mM sodium phosphate buffer, pH 4.5, containing 300 mM NaCl (Buffer B). After nonabsorbed substances had been washed out of the column with Buffer B, the recombinant Vag8 proteins were eluted with Buffer B containing 300 mM imidazole. Imidazole in the Vag8 fraction was removed by dialysis against Buffer A. Protein concentrations were determined using a Micro BCA Protein Assay Kit (Thermo Fisher Scientific). The recombinant proteins were subjected to SDS-PAGE followed by immunoblotting using anti-HAT-tag antibody (GenScript) and HRP-conjugated anti-rabbit IgG (Jackson ImmunoResearch). Target proteins were visualized with Immobilon Western Chemiluminescent HRP Substrate (Merck Millipore).

### (ii) ELISA-based binding assay

Wells of 96-well microplates (ELISA Plate H; Sumitomo Bakelite) were coated with 0.1 ml of 10 nM C1 Inh (Complement C1 Inhibitor, Human; Calbiochem) at 4°C overnight, and then blocked with Dulbecco’s–modified phosphate buffered saline (D-PBS) containing 10% skim milk at 37°C for 1 hour. Recombinant Vag8 proteins at the indicated concentrations were added to the wells and allowed to react at 37°C for 1 hour. The Vag8 proteins bound to C1 Inh were probed with a combination of anti-HAT-tag antibody, HRP-conjugated anti-rabbit IgG, and 0.1% TMBZ (Dojindo Laboratories). Each antibody reaction was carried out at 37°C for 1 hour. The substrate reaction was carried out at room temperature for 30 minutes and stopped by addition of 1.0 N H_2_SO_4_. After each step before the substrate reaction, the wells were washed with D-PBS containing 0.05% Tween-20. The optical density at 450 nm (OD_450_) of each well was read with a Multi-Detection Microplate Reader (POWERSCAN HT; BioTek).

### (iii) Inhibitory effects of C1 Inh on PK activity

Recombinant Vag8 proteins (1 µM) were incubated with 50 nM C1 Inh, and subsequently with 2 nM PK (Human Kallikrein; Enzyme Research Laboratories) at 37°C for 1 hour each. After incubation, the solution was mixed with the chromogenic substrate (S-2302, 0.5 mM; Chromogenix) in HEPES-NaHCO_3_ buffer (9), and incubated at 37°C for 1 hour. The OD_405_ value of each well, determined using a Multi-Detection Microplate Reader, was referred to as the protease activity of PK.

### (iv) PK-C1 Inh complex formation assay

Recombinant Vag8 proteins (3 µM) were preincubated with 100 nM C1 Inh at 37°C for 1 hour in 27 mM HEPES buffer, pH 7.4, containing 191 mM NaCl and 0.68 mM EDTA (10). The resultant mixture was mixed with 100 nM PK, incubated at 37°C for 1 hour, and subjected to SDS-PAGE followed by immunoblotting using anti-C1 Inh antibody (Abcam) and HRP-conjugated anti-rabbit IgG as described above.

## Acknowledgements

This work was supported by JSPS KAKENHI Grant Numbers 17H04075 and 19K16638.

## References

1. Tan T, Dalby T, Forsyth K, Halperin SA, Heininger U, Hozbor D, Plotkin S, Ulloa-Gutierrez R, Wirsing von König CH. 2015. Pertussis across the globe: recent epidemiologic trends from 2000 to 2013. Pediatr Infect Dis J 34:e222–32.

2. Mills KHG, Ross PJ, Allen AC, Wilk MM. 2014. Do we need a new vaccine to control the re-emergence of pertussis? Trends Microbiol 22:49–52.

3. Suzuki K, Shinzawa N, Ishigaki K, Nakamura K, Abe H, Fukui-Miyazaki A, Ikuta K, Horiguchi Y. 2017. Protective effects of *in vivo*-expressed autotransporters against *Bordetella pertussis* infection. Microbiol Immunol 61:371–379.

4. de Gouw D, Gouw D de, de Jonge MI, Jonge MI de, Hermans PWM, Wessels HJCT, Zomer A, Berends A, Pratt C, Berbers GA, Mooi FR, Diavatopoulos DA. 2014. Proteomics-identified Bvg-activated autotransporters protect against *Bordetella pertussis* in a mouse model. PLoS ONE 9:e105011.

5. Hovingh ES, van den Broek B, Kuipers B, Pinelli E, Rooijakkers SHM, Jongerius I. 2017. Acquisition of C1 inhibitor by *Bordetella pertussis* virulence associated gene 8 results in C2 and C4 consumption away from the bacterial surface. PLoS Pathog 13:e1006531.

6. Davis AE, Mejia P, Lu F. 2008. Biological activities of C1 inhibitor. Mol Immunol 45:4057–4063.

7. Hovingh ES, de Maat S, Cloherty APM, Johnson S, Pinelli E, Maas C, Jongerius I. 2018. Virulence associated gene 8 of *Bordetella pertussis* enhances contact system activity by inhibiting the regulatory function of complement regulator C1 inhibitor. Front Immunol 9:1172.

8. Marr N, Shah NR, Lee R, Kim EJ, Fernandez RC. 2011. Bordetella pertussis autotransporter Vag8 binds human C1 esterase inhibitor and confers serum resistance. PLoS ONE 6:e20585.

9. Kolte D, Bryant J, Holsworth D, Wang J, Akbari P, Gibson G, Shariat-Madar Z. 2011. Biochemical characterization of a novel high-affinity and specific plasma kallikrein inhibitor. Br J Pharmacol 162:1639–1649.

10. Beinrohr L, Harmat V, Dobó J, Lörincz Z, Gál P, Závodszky P. 2007. C1 inhibitor serpin domain structure reveals the likely mechanism of heparin potentiation and conformational disease. J Biol Chem 282:21100–21109.

