## Supplementary material for "Localization of regions of a *Bordetella pertussis* autotransporter, Vag8, interacting with C1 inhibitor": Table S1

**Table S1.** Primers used in this study

| Primers | Sequence (5’-3’) | Vag8 derivatives |
| --- | --- | --- |
| vag8-*Xho*I | CACAACAAGCTCGAGTTTTCTGGGCGATGTC | Vag8_WT_ |
| vag8-*Eco*RI | AAGCTTGAATTCCTACCCGATCGAATGCAG | Vag8_WT_ |
| pVag1 | ATGCCCACAACAAGCTCGAGTTTCTGG | Vag8_52-596, 52-548, 52-479_ |
| pVag2 | CTGCAGGTCGACAAGCTCTACCGGGAATCGCCCGTCACG | Vag8_52-479, 102-479_ |
| pVag6 | ACTGCAGGTCGACAAGCTTGAATTCC | Vag8_102-648, 152-648, 202-648_ |
| pVag9 | TGCCCACAACAAGCTCGAGCCGGACAACACCGCGCT | Vag8_102-648, 102-596, 102-548, 102-479_ |
| pVag10 | TGCCCACAACAAGCTCGAGGGCGATGACAGTTTCGCCCT | Vag8_152-648_ |
| pVag11 | TGCCCACAACAAGCTCGAGGGGGCAGGCGTTTCCGCGA | Vag8_202-648_ |
| pVag14 | CTGCAGGTCGACAAGCTCTACGTGTTGTTGACCACCAGCA | Vag8_52-548, 102-548_ |
| pVag15 | CTGCAGGTCGACAAGCTCTAGGCCTGCGCCTCGC | Vag8_52-596, 102-596_ |
